# Multimodal MR Imaging for quantification of brain lipid in mice at 9.4T

**DOI:** 10.1101/2025.09.30.679612

**Authors:** Sunil Kumar Khokhar, Anshuman Swain, Narayan Datt Soni, Halvor Juul, Abeer Mathur, Dipak Roy, Blake Benyard, Dushyant Kumar, Ravi Prakash Reddy Nanga, Mohammad Haris, Ravinder Reddy

**Affiliations:** Center for Advanced Metabolic Imaging in Precision Medicine, Department of Radiology, University of Pennsylvania, Philadelphia, PA, United States; School of Engineering and Applied Science, University of Pennsylvania, Philadelphia, PA, United States

**Author notes:** Corresponding Author Prof. Ravinder Reddy, Director, Center for Advanced Metabolic Imaging in Precision Medicine, Department of Radiology, University of Pennsylvania, Philadelphia, PA, United States. Share equal authorship (Sunil Kumar Khokhar, Anshuman Swain).

**Keywords:** Brain, Lipid Imaging, Myelination, Repeatability, NOE_MTR_, tNOE, MWF

## Abstract

**Background:** Advanced MR imaging techniques like steady state Nuclear Overhauser enhancement (ssNOE), transient NOE (tNOE), and myelin water fraction (MWF) provide a non-invasive way to assess the biochemical and structural integrity of brain tissue. Their sensitivity to endogenous lipids and macromolecules allows for the early detection of neuropathological changes, making them valuable tools in studying brain health and disease progression. In this study, we systematically evaluate the repeatability and sensitivity of NOE_MTR_, tNOE, and MWF for quantifying lipid and myelin content in the brains of wild-type (WT) mice, correlating the results with immunohistochemistry (IHC).

**Methods:** Five 6-month-old C57BL6/J mice were imaged using 3D-NOE, and four mice underwent imaging with 2D tNOE and MWF across four repeated sessions using a 9.4T Scanner. For ssNOE imaging, CEST-weighted images at 56 frequency offsets were acquired using B_1rms_ of 1.0 μT and 3s saturation duration. For tNOE, 52 offsets were acquired with a hyperbolic secant inversion pulse (bandwidth = 400Hz, duration = 44ms) and a mixing time of 200ms. For MWF, a multi-echo spin-echo (MESE) sequence was acquired with 40 evenly spaced echoes from 5.5ms to 200ms. For both ssNOE and tNOE, B_0_ correction was performed using WASSR. Repeatability was quantified using intra- and inter-subject coefficients of variation (COV%). Pearson correlation was performed to see the association between imaging matrices and IHC measures, Luxol fast blue (LFB) stained sections, and myelin basic protein (MBP).

**Results:** All techniques demonstrated high repeatability across the whole brain (WB) and selected regions of interest (ROIs). Whole-brain intra-subject COV% for NOE_MTR_ ranged from 1.92% to 3.40%, with corresponding inter-subject COVs of 1.50%. tNOE exhibited improved intra-subject repeatability with COVs ranging from 0.75% to 5.57%, but a reduced inter-subject COV of 2.97%. MWF imaging showed the highest stability overall, with an intra-subject COV ranging from 0.47% to 2.03% and an inter-subject COV of 0.75%. Visually, tNOE offers superior contrast in myelin-rich areas compared to NOE_MTR_ and MWF imaging, showing greater sensitivity to myelinated regions. tNOE strongly correlates with histological markers: r = 0.83 with MBP staining and r = 0.72 with LFB staining (both p < 0.001). MWF and NOE_MTR_ showed correlations with MBP (r = 0.63 and r = 0.57, respectively).

**Conclusion:** NOE_MTR_, tNOE, and MWF imaging are reliable and repeatable methods for quantifying macromolecules in the brain. Among these, tNOE emerges as the most sensitive for detecting myelin lipids as confirmed by histological validation. These findings highlight the translational potential of tNOE for studying demyelinating disorders and neurodegenerative diseases.

## BACKGROUND

Lipids are essential to brain function, serving as structural components of cell membranes and the myelin sheath that insulates nerve fibers and enables rapid signal transmission (1). Beyond these structural roles, lipids contribute critically to brain development, especially during infancy, and play a key role in supporting cognitive functions such as learning and memory. They also act as energy sources during times of glucose shortage and participate in important signaling pathways that regulate neuronal growth, communication, and repair (2). Disruptions in lipid homeostasis have been associated with neurological and neurodegenerative conditions such as multiple sclerosis, Alzheimer’s disease, and depression (3). Specifically, myelin lipids in the central nervous system are critical for preserving signal conduction and neuronal integrity (4), and their breakdown is a hallmark of several neurodegenerative disorders (5,6). As such, non-invasive imaging techniques that quantify brain lipid content offer valuable tools for early diagnosis, disease monitoring, and evaluating treatments. Magnetic resonance (MR) imaging has transformed the non-invasive examination of the brain by providing detailed insights into its structural, functional, and molecular characteristics. Recent advances in MRI techniques, such as steady state Nuclear Overhauser Enhancement (ssNOE), transient NOE (tNOE), and myelin water imaging, have expanded the ability to detect macromolecular changes, including lipids in the brain (7–10). These methods are being increasingly applied in neuroscience research, particularly in studying neurodegenerative diseases, demyelinating disorders, and other brain pathologies (7,11,12).

The Nuclear Overhausser Effect (NOE, referred to as steady state NOR or ssNOE), was originally introduced by Albert Overhauser to describe polarization transfer of spins in metals (13). Since then, it has become a cornerstone technique in high-resolution NMR for probing the conformational structure of macromolecules. Previously, ssNOE has been adapted for *in vivo* neuroimaging (14), with the effect resulting from dipolar interactions between macromolecules, such as lipids and proteins, and bulk water in the brain, making it highly relevant for studying myelin integrity and cellular membranes (15,16). Metrics such as NOE Magnetization Transfer Ratio (NOE_MTR_) and relayed NOE (rNOE) components have been presented for comprehensive characterization of brain tissues (7,17). In addition, ssNOE imaging has been used to evaluate demyelinating conditions, such as multiple sclerosis, where decreased ssNOE signals correlate with the loss of myelin-associated macromolecules (18). However, traditional steady-state NOE imaging can be time-consuming as it requires the acquisition of the canonical Z-spectra followed by multi-pool fitting to derive the aforementioned metrics. In addition, steady-state NOE metrics can be heavily biased by confounding factors such as direct water saturation, magnetization transfer effects, and field inhomogeneities (B_0_, B_1_). To address these limitations, tNOE imaging has been developed, offering a more time-efficient approach with reduced MT and spillover effects. By employing a frequency-selective inversion pulse followed by a mixing time, in which cross-relaxation from aliphatic protons of lipids and macromolecules to bulk water occurs, tNOE can capture information from lipid-rich environments with reduced MT and direction saturation contributions. Recent studies have demonstrated the feasibility of tNOE in human brain imaging at 7T, highlighting its potential for clinical applications (9).

Traditionally, myelin water imaging (19) has been utilized to extract a quantitative measure of myelin content, known as myelin water fraction (MWF), by analyzing the multi-exponential T_2_ relaxation of water protons (20). The short T_2_ component, attributed to water trapped between myelin bilayers, serves as a surrogate marker for myelin density (21). MWF has been validated against histological measures and has shown high reproducibility across different imaging protocols and MRI scanners (4). Previous studies have validated MWF as a reliable imaging biomarker in both preclinical and clinical settings, particularly for disorders characterized by white matter pathology (22,23). However, MWF measures the myelin-associated water and does not provide a direct measure of myelin lipids, which may preclude applications to pathologies that exhibit diffuse lipid changes prior to lipid degeneration and corresponding decrease in the myelin water fraction.

A comprehensive understanding of these competing quantitative imaging techniques’ strengths and limitations is essential for optimizing their use in brain lipid research across both preclinical and clinical settings. In this study, we employ wild-type mouse brains to systematically evaluate the repeatability and sensitivity of NOE_MTR_, tNOE, and myelin water imaging for quantifying lipid and myelin content. We further validate these imaging biomarkers through immunohistochemistry (IHC) for lipid and myelin. Our aim is to elucidate the capabilities of these techniques in detecting myelin-rich regions and to explore their potential applications in neurodegenerative disease research.

## METHODS

### Animal Preparation

We performed 3D steady-state NOE imaging on five 6-month-old C57BL6/J mice (3 females and 2 males), and 2D tNOE and MWF on four additional mice (2 females and 2 males). All experimental procedures were approved by the Institutional Animal Care and Use Committee (IACUC) at the University of Pennsylvania under protocol number (IACUC/807508/2024). The mice were anesthetized with 1.5% isoflurane, which was delivered continuously in an oxygen mixture. Their heads were secured in a conical head restrainer, and a respiratory monitoring pad, along with a rectal temperature probe, was used to continuously track their breathing rate and body temperature (24). Throughout the experiment, the respiration rate and body temperature were maintained at 60–90 breaths per minute and 37°C, respectively. The animal was positioned inside a 20 mm dual-channel ^1^H transceiver volume coil, ensuring that the brain region of interest was aligned at the center of the coil. This assembly was then placed inside a 30 cm horizontal-bore 9.4 T magnet, which was interfaced with an Avance III HD console and Paravision 6.0.1 software (Bruker BioSpin, Germany). Each animal underwent four scans within a week to verify the repeatability of the scan results (Fig. 1).

**Figure 1:**
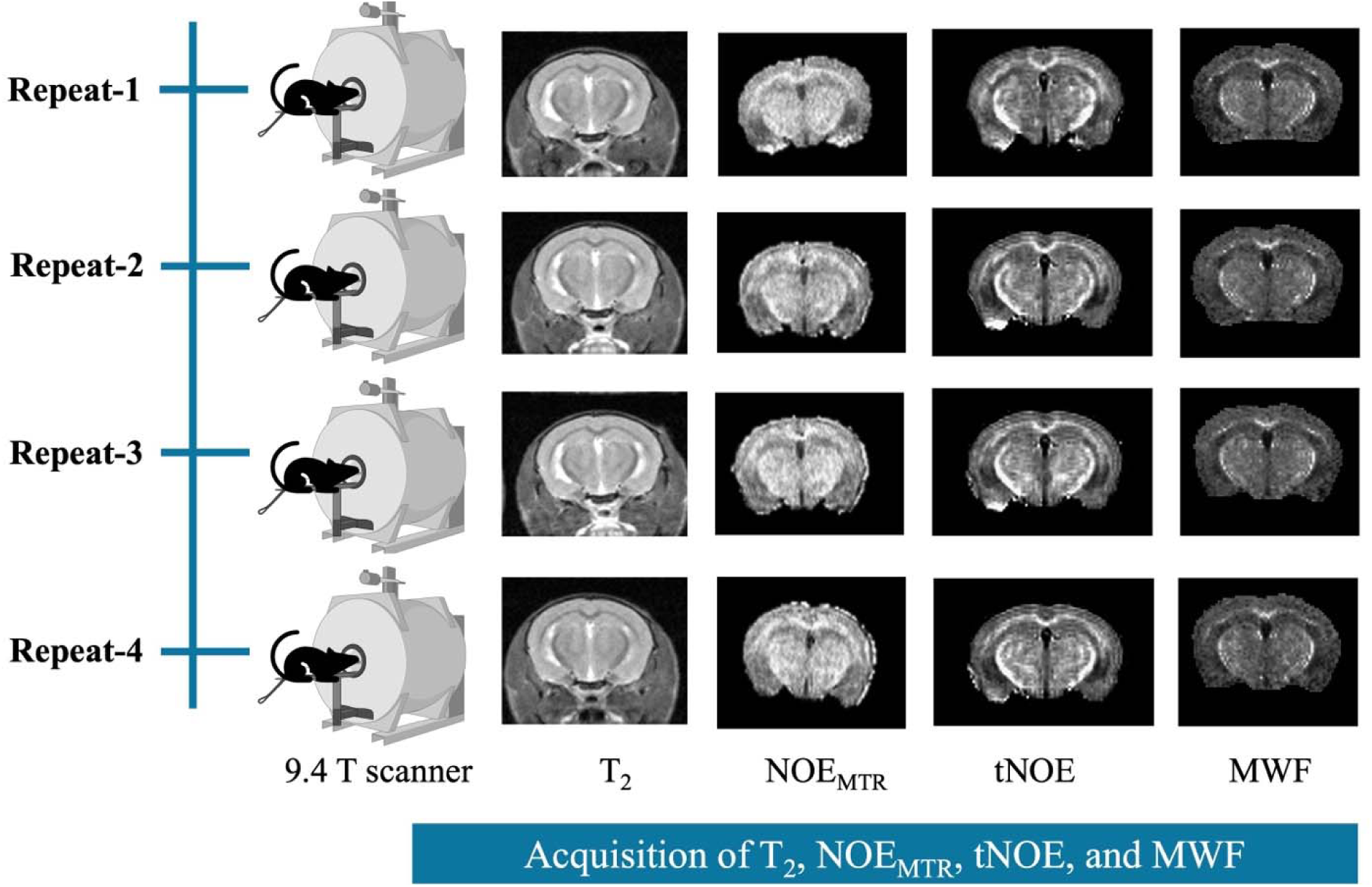
Schematic representation of the repeated imaging sessions (Repeat-1 to Repeat-4) performed on the same animal using a preclinical MR imaging scanner. Each session included the acquisition of advanced MR imaging sequences: NOE, transient NOE (tNOE), and myelin water fraction (MWF) imaging. From these scans, contrast metrics were derived, including NOE_MTR_, tNOE, and MWF.

### MRI acquisition

A localizer scan was initially acquired to ensure proper positioning of the animal. Once the correct alignment was achieved, the frequency for both channels was tuned to the ^1^H resonance, and the global frequency was adjusted accordingly. Automated global shimming of both first- and second-order shims was then performed to achieve a smooth and symmetric water signal. The reference power was calibrated using a 5mm slice positioned around the isocenter, and a B_0_ map was acquired for subsequent shimming. T_2_-weighted rapid acquisition with relaxation enhancement (RARE) images were acquired with the following parameters: repetition time (TR) = 1500 ms, echo time (TE) = 12.24 ms, 2 averages, 16 axial slices of 1 mm thickness, RARE factor = 4, field of view (FOV) = 12 × 15 mm², and a matrix size = 80 × 100. Both 2D and 3D T_2_-weighted images were used for structural reference and registration to a mouse brain template.

### 3D ssNOE Imaging

Steady state NOE data were acquired using a magnetization-preparation module consisting of a train of 15 rectangular saturation pulses (B_1, RMS_ = 1.0 μT, duration = 3.0 s each, no inter-pulse delay, continuous wave), followed by multi-shot (3 segments), 3D GRE-based readout. For acquiring corresponding spectra, ssNOE data were acquired with frequency offsets ranging from 100 ppm to -100 ppm (Supplementary Table 1), with denser sampling between 0 and 5 ppm and sparser sampling elsewhere.

Additional acquisition parameters were: averages = 2, number of frequency offsets = 56, Flip Angle = 5^°^, TE = 2 ms, shot TR= 6.0 s, FOV = 12 × 15 x 10 mm^3^, matrix size = 80 × 100 x 10, and partial-Fourier = 70%. Images were reconstructed using projection onto convex sets (POCS) to recover the missing k-space data due to partial Fourier acquisitions (25). For correction of B_0_ inhomogeneity, a B_0_ map was generated using a Water Saturation Shift Referencing (WASSR) (26) image with the following parameters: TR/TE = 2512.4/2.10 ms, with 22 frequency offsets ranging from 0 to ±1 ppm at a step size of 0.1 ppm, B_1rms_ = 0.1 μT, FOV = 12 × 15 mm², matrix size: 80 × 100, and a single average. Additionally, an unsaturated image was acquired using the same parameters as the NOE-weighted images with an offset of -300ppm.

### 2D tNOE MR Imaging

Transient NOE data were acquired using a magnetization-preparation module consisting of a frequency-selective hyperbolic secant inversion pulse (duration 44 ms, inversion bandwidth = 400 Hz), followed by a variable mixing-time delay and then a single-shot GRE-based readout. For acquiring corresponding spectra, tNOE data were acquired with frequency offsets ranging from 5 ppm to -5 ppm (Supplementary Table 2). Additional acquisition parameters were: number of averages = 3, number of offsets = 52, Flip Angle = 10°, TE = 3 ms, shot TR = 8.0 s, FOV = 12 × 15 mm², matrix size = 80 × 100, 1mm slice thickness, and mixing time of 200 ms. WASSR was employed for B_0_ inhomogeneity correction using the same preparation as the ssNOE imaging, with changes to the following parameters: -TR/TE = 410.4/4 ms and a single slice of 1mm thickness. Also, an unsaturated image was acquired using the same parameters as the tNOE-weighted images with an offset of -300ppm. We have applied a saturation pulse at -3.5 ppm relative to the water resonance to probe tNOE contributions, mainly arising from aliphatic groups in lipids and proteins.

### Myelin Water Imaging

A multi-echo spin echo (MESE) sequence was used to determine the MWF following non-negative least squares (NNLS) fitting. The acquisition parameters were as follows: TR = 6400 ms, number of echoes = 40 with echo times ranging from 5.5 ms to 220 ms, resulting in an echo spacing of (ΔTE) = 5.5 ms. The FOV was set to 20 mm × 20 mm with an acquisition matrix of 128 × 128. A total of 8 contiguous slices were acquired, each with a slice thickness of 1 mm and no inter-slice gap, and the number of averages = 2.

### Image processing and analysis

Image processing was conducted using MATLAB 2023b (MathWorks, CA). T_2_-weighted (T_2w_) images were subjected to automated skull stripping using SHERM (27), with errors in the brain mask corrected using ITK-SNAP (28). For image registration, we utilized a C57BL/6 J mouse brain atlas provided by (29) as a template. This template comprised a three-dimensional T_2w_ RARE image and detailed brain parcellations covering the entire mouse brain. For 3D image registration, a custom low-resolution atlas was generated using the average of T_2w_ images acquired from each mouse registered to the same reference space. The Dorr atlas was then registered to this custom low-resolution atlas using ANTS (30), with the transformation matrix applied to the parcellations using nearest-neighbor interpolation. The custom atlas was then registered to each mouse brain to get parcellations specific to each mouse with their respective repeated acquisitions. For 2D image registration, the corresponding slice from the T_2w_ image acquired was adjusted to match the intensity of the atlas image through histogram-based normalization (31). To facilitate registration, both the corresponding slice from the Dorr atlas and target images were incrementally blurred with a Gaussian filter, then aligned using an affine transformation across ten iterations, with the Gaussian kernel’s width gradually reduced to improve the matching of finer anatomical structures. Non-linear registration was subsequently performed using the Demons algorithm (32). The corresponding segmentations from the atlas were aligned by applying the transformations to the atlas labels using nearest-neighbor interpolation. NOE_MTR_ images were generated using the following equation (12):

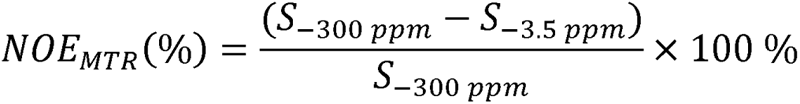

where S_-3.5ppm_ and S_-300ppm_ denote the signal magnitudes from an image voxel at offsets of _-3.5 ppm_ and _-300 ppm_, respectively. NOE_MTR_ was corrected for B_0_ inhomogeneities.

To quantify MWF, we employed the MWF model available in the qMRLab toolbox (https://qmrlab.org/). The analysis was based on spin-echo images acquired at 40 echo times ranging from 5.5 ms to 220 ms, enabling robust sampling of T_2_ decay curves. Skull-stripping was performed on the T_2w_ image as described earlier to generate the brain mask and restrict the voxel-wise fitting to brain tissue and enhance computational efficiency. The model was initialized, and the data and mask were loaded into a structure and fitted using the FitData function, which estimated voxel-wise maps of MWF. The MWF is calculated as the fraction of signal with short T_2_ values (e.g., <40 ms), representing myelin water, relative to the total water fraction. This method enables non-invasive, compartment-specific mapping of myelin content in brain tissue (33).

We quantified the contrast values for NOE_MTR_ and MWF from the whole brain (WB) and for tNOE for whole slice and subregions comprising the cerebral cortex, entorhinal cortex, hippocampus, thalamus, hypothalamus, corpus callosum, cerebral peduncle, and fimbria. Mean contrast values for NOE_MTR_, tNOE, and MWF from these subregions were calculated using the registered template parcellations.

### Immunohistochemistry

All histopathology and immunohistochemistry were performed at the core facility of the Penn Center for Musculoskeletal Disorders, at the University of Pennsylvania. Mouse brains were fixed through transcardiac perfusion and removed, followed by a post-fixation in 4% paraformaldehyde. The brain was sliced into 2mm coronal blocks and embedded in a paraffin block. The specimen was sectioned at 10 µm sections, and serial sections were collected and mounted on glass slides.

For Luxol fast blue (LFB) staining, formalin-fixed paraffin-embedded (FFPE) brain sections were baked at 60°C for 30 minutes to melt the paraffin, followed by deparaffinization through sequential incubations in xylene (3 × 5 min), 100% ethanol (2 × 3 min), and 95% ethanol (2 × 5 min). Sections were then incubated overnight at 60°C in 0.1% LFB solution prepared in 95% ethanol with acetic acid. After rinsing in double-distilled water (ddH O), differentiation was performed using 0.05% lithium carbonate solution for up to 3 minutes, followed by 70% ethanol until a clear contrast between white and gray matter was observed. Differentiation was assessed under a microscope and repeated if necessary. Final dehydration was carried out in 95% ethanol, 100% ethanol (2 × 2 min), and xylene (2 × 2 min), and slides were mounted using Permount.

For IHC detection of myelin basic protein (MBP), FFPE sections were baked at 60°C for 15-20 minutes and rehydrated through a graded series of xylene (3 × 5 min), 100% ethanol (2 × 3 min), 95% ethanol (2 min), and 70% ethanol (2 min), followed by a brief rinse in ddH O. Antigen retrieval was performed by heating the sections in 1× Tris-EDTA buffer (pH 8.0) in a pressure cooker for 15 minutes, followed by cooling for 10 minutes. Slides were washed in 1× PBS and incubated in 3% hydrogen peroxide for 15 minutes to block endogenous peroxidase activity. After PBS-T washes (3 × 5 min), a hydrophobic barrier was drawn around each section, and slides were blocked with 3% BSA in PBS for 1 hour. Primary anti-MBP antibody was applied and incubated overnight in a humidified chamber. The next day, sections were washed and incubated with biotinylated secondary antibodies. DAB substrate was applied until optimal staining intensity was achieved. Sections were counterstained, dehydrated in graded alcohols and xylene.

After staining, slides were scanned using a high-resolution digital slide scanner. Quantification of stain intensity was performed using FIJI (ImageJ), where staining intensity was measured from manually defined ROIs in white matter and gray matter areas, including the corpus callosum, fimbria, cerebral peduncle, hippocampus, and cerebral cortex.

### Statistical analysis

The coefficient of variation (COV) was quantified to evaluate the variability of contrast both within and between subjects across multiple imaging sessions (34). It is calculated as the ratio of the standard deviation (σ) to the mean (μ) of contrast values.

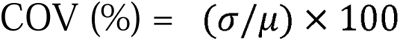

For intra-subject analysis, the COV was calculated by dividing the standard deviation of mean contrast values from the whole brain and sub-regions by the means of these regions. For inter-subject analysis, the mean contrast value was calculated across all repeat scans for each subject (whole-brain and sub-regions). The inter-subject COV was calculated by taking the standard deviation of these means and dividing it by their mean. Pearson’s correlation coefficients were calculated (scipy.stats, Python) to evaluate the relationship between imaging metrics and immunohistochemical outcomes,

## RESULTS

The repeatability results for NOE_MTR_ (Fig. 2), MWF (Fig. 3), and tNOE (Fig. 4) were evaluated across various brain regions, including the whole brain, hippocampus, cerebral cortex, thalamus, hypothalamus, corpus callosum, fimbria, and cerebral peduncle. The COVs measured for WB and various ROIs for different imaging methods are shown in Tables 1 and 2.

**Figure 2:**
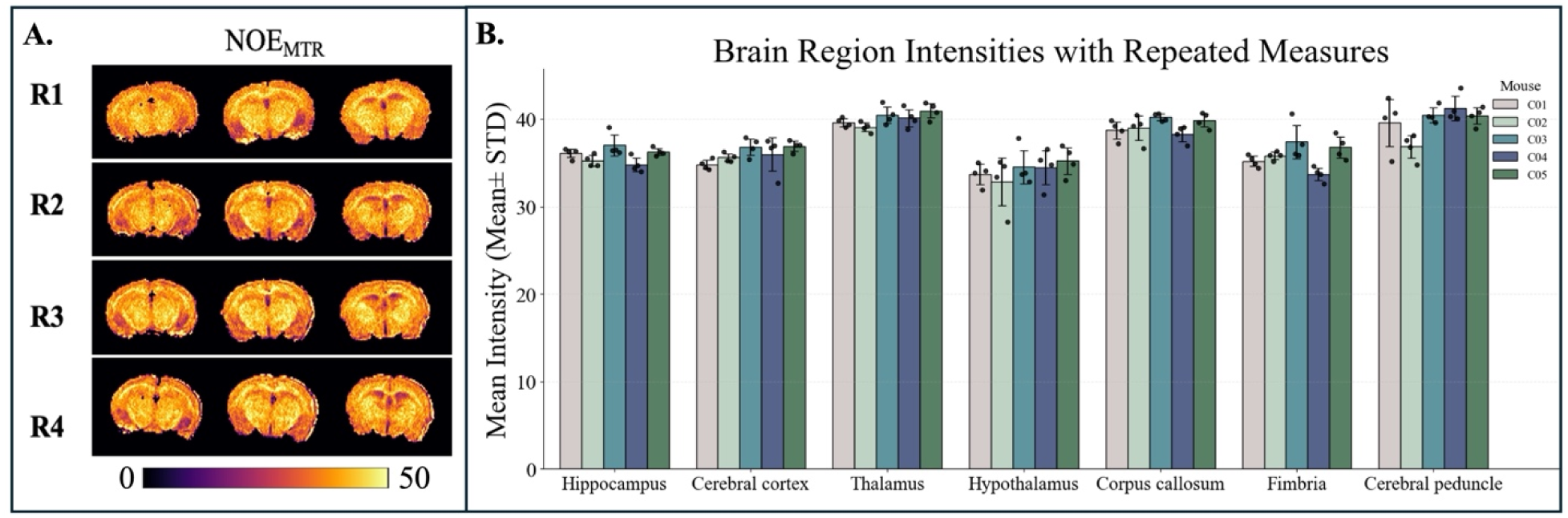
**A)** Representative whole brain global maps from a WT mouse with four repeats (R1-R4) of NOE_MTR_, map contrast, B) Bar graph showing the mean NOE_MTR_ intensities (mean ± standard deviation) across seven brain regions for five WT mice (C1–C5), each with four repeated measures.

**Table 1:** The intra- and inter-subject COV% for NOE_MTR_, tNOE, and MWF map contrast from the whole brain.

| Mouse numbers |  | NOE<sub>MTR</sub> | tNOE | MWF |
| --- | --- | --- | --- | --- |
| **C1** | **Mean** | 36.25 | 8.00 | 56.79 |
|  | **STD** | ±0.91 | ±0.10 | ±1.03 |
|  | **COV%** | **2.50** | **1.26** | **1.82** |
| **C2** | **Mean** | 36.06 | 8.39 | 57.00 |
|  | **STD** | ±0.75 | ±0.47 | ±0.27 |
|  | **COV%** | **2.08** | **5.57** | **0.47** |
| **C3** | **Mean** | 37.19 | 7.98 | 60.34 |
|  | **STD** | ±1.14 | ±0.29 | ±0.49 |
|  | **COV%** | **3.05** | **3.63** | **0.81** |
| **C4** | **Mean** | 36.34 | 7.83 | 58.74 |
|  | **STD** | ±1.24 | ±0.06 | ±1.19 |
|  | **COV%** | **3.40** | **0.75** | **2.03** |
| **C5** | **Mean** | 37.03 | NA | NA |
|  | **STD** | ±0.71 | NA | NA |
|  | **COV%** | **1.92** | **NA** | **NA** |
| **Inter-subject** | **Mean** | 36.40 | 8.05 | 58.22 |
|  | **STD** | ±0.55 | ±0.24 | ±0.44 |
|  | **COV%** | **1.50** | **2.97** | **0.75** |

**Table 2:**
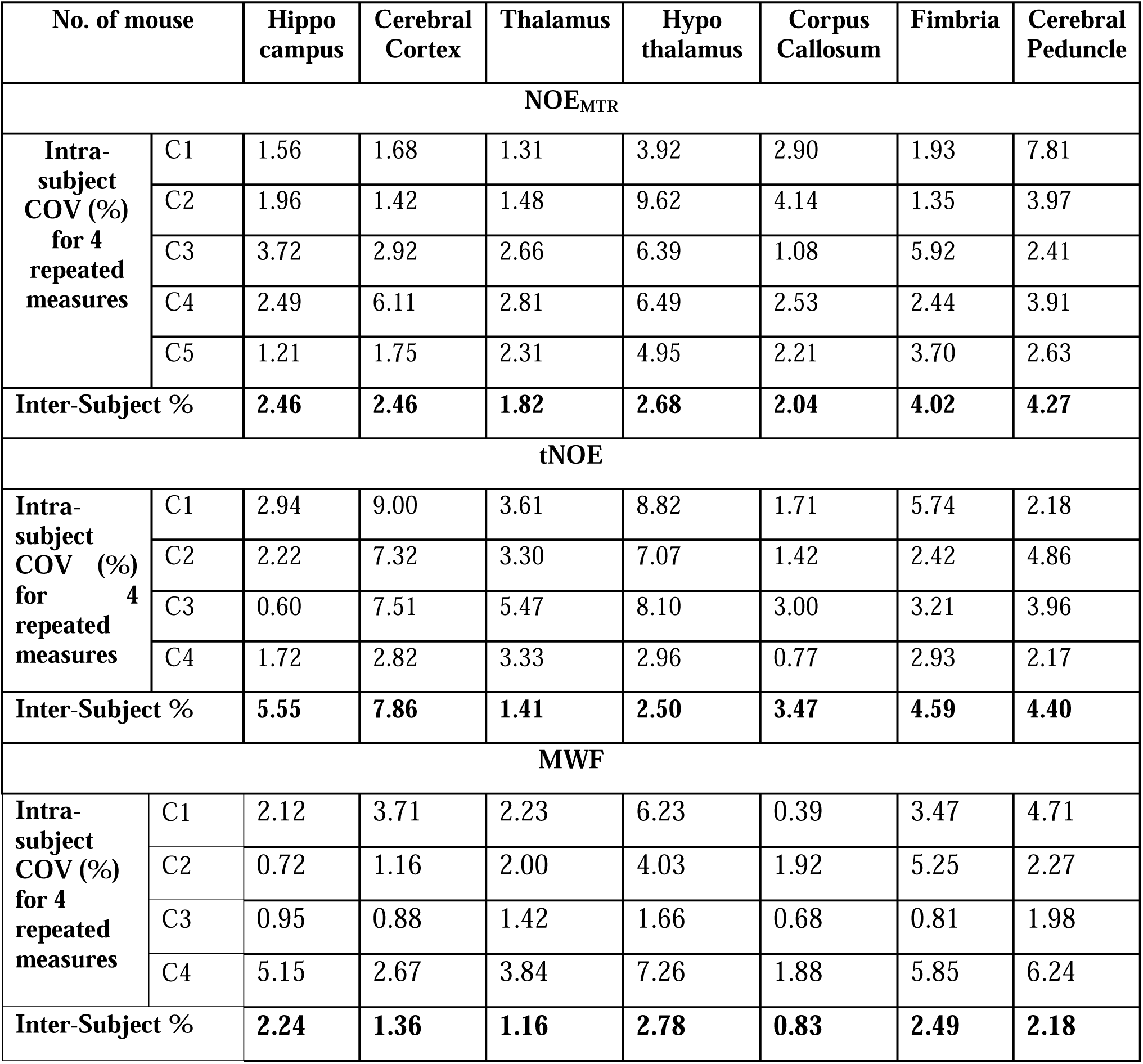
The intra- and inter-subject COV% of each ROI for matrices of NOE_MTR_, tNOE, and MWF map contrast.

| No. of mouse |  | Hippo<br>campus | Cerebral<br>Cortex | Thalamus | Hypo<br>thalamus | Corpus<br>Callosum | Fimbria | Cerebral<br>Peduncle |
| --- | --- | --- | --- | --- | --- | --- | --- | --- |
| **NOE<sub>MTR</sub>** |  |  |  |  |  |  |  |  |
| **Intra-**<br>**subject**<br>**COV (%)**<br>**for 4**<br>**repeated**<br>**measures** | C1 | 1.56 | 1.68 | 1.31 | 3.92 | 2.90 | 1.93 | 7.81 |
|  | C2 | 1.96 | 1.42 | 1.48 | 9.62 | 4.14 | 1.35 | 3.97 |
|  | C3 | 3.72 | 2.92 | 2.66 | 6.39 | 1.08 | 5.92 | 2.41 |
|  | C4 | 2.49 | 6.11 | 2.81 | 6.49 | 2.53 | 2.44 | 3.91 |
|  | C5 | 1.21 | 1.75 | 2.31 | 4.95 | 2.21 | 3.70 | 2.63 |
| **Inter-Subject %** |  | **2.46** | **2.46** | **1.82** | **2.68** | **2.04** | **4.02** | **4.27** |
| **tNOE** |  |  |  |  |  |  |  |  |
| **Intra-**<br>**subject**<br>**COV (%)**<br>**for 4**<br>**repeated**<br>**measures** | C1 | 2.94 | 9.00 | 3.61 | 8.82 | 1.71 | 5.74 | 2.18 |
|  | C2 | 2.22 | 7.32 | 3.30 | 7.07 | 1.42 | 2.42 | 4.86 |
|  | C3 | 0.60 | 7.51 | 5.47 | 8.10 | 3.00 | 3.21 | 3.96 |
|  | C4 | 1.72 | 2.82 | 3.33 | 2.96 | 0.77 | 2.93 | 2.17 |
| **Inter-Subject %** |  | **5.55** | **7.86** | **1.41** | **2.50** | **3.47** | **4.59** | **4.40** |
| **MWF** |  |  |  |  |  |  |  |  |
| **Intra-**<br>**subject**<br>**COV (%)**<br>**for 4**<br>**repeated**<br>**measures** | C1 | 2.12 | 3.71 | 2.23 | 6.23 | 0.39 | 3.47 | 4.71 |
|  | C2 | 0.72 | 1.16 | 2.00 | 4.03 | 1.92 | 5.25 | 2.27 |
|  | C3 | 0.95 | 0.88 | 1.42 | 1.66 | 0.68 | 0.81 | 1.98 |
|  | C4 | 5.15 | 2.67 | 3.84 | 7.26 | 1.88 | 5.85 | 6.24 |
| **Inter-Subject %** |  | **2.24** | **1.36** | **1.16** | **2.78** | **0.83** | **2.49** | **2.18** |

**Figure 3:**
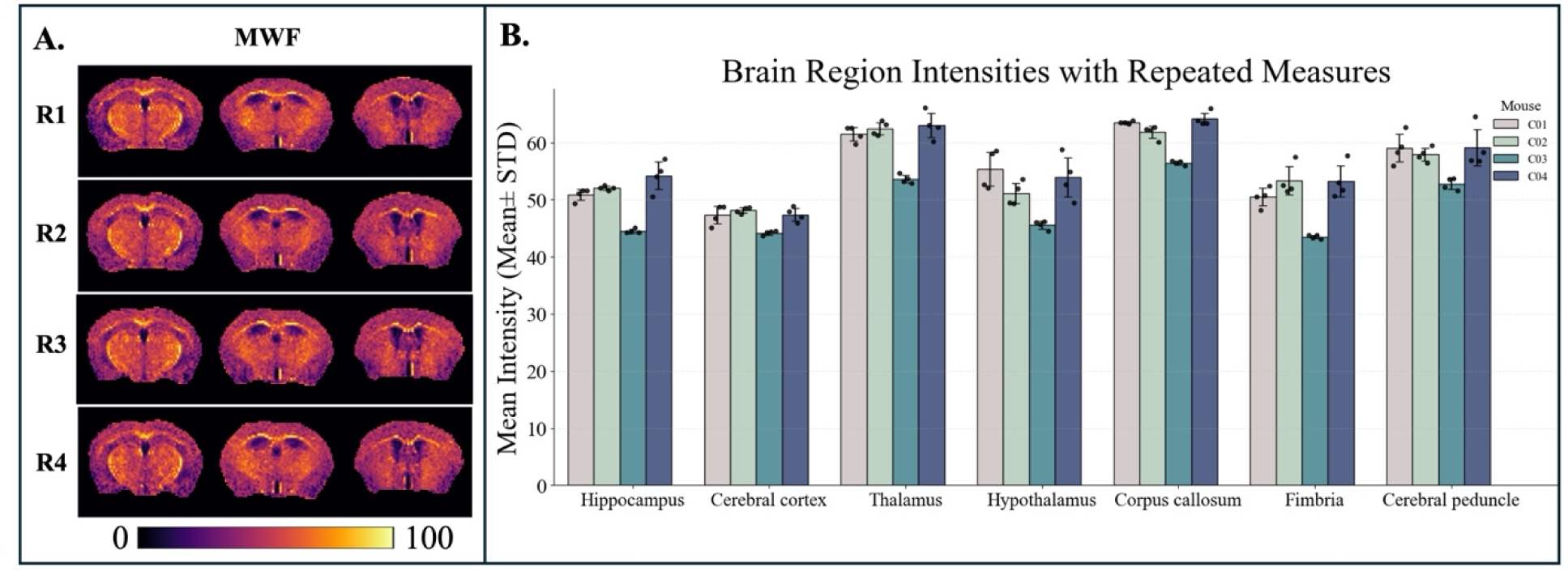
A) Representative coronal slices from four repeated sessions (R1-R4) show consistent spatial patterns in myelin water fraction (MWF) maps. B) Bar graph showing the mean MWF intensities (mean ± standard deviation) across seven brain regions for four WT mice (C1–C4), each with four repeated measures.

**Figure 4:**
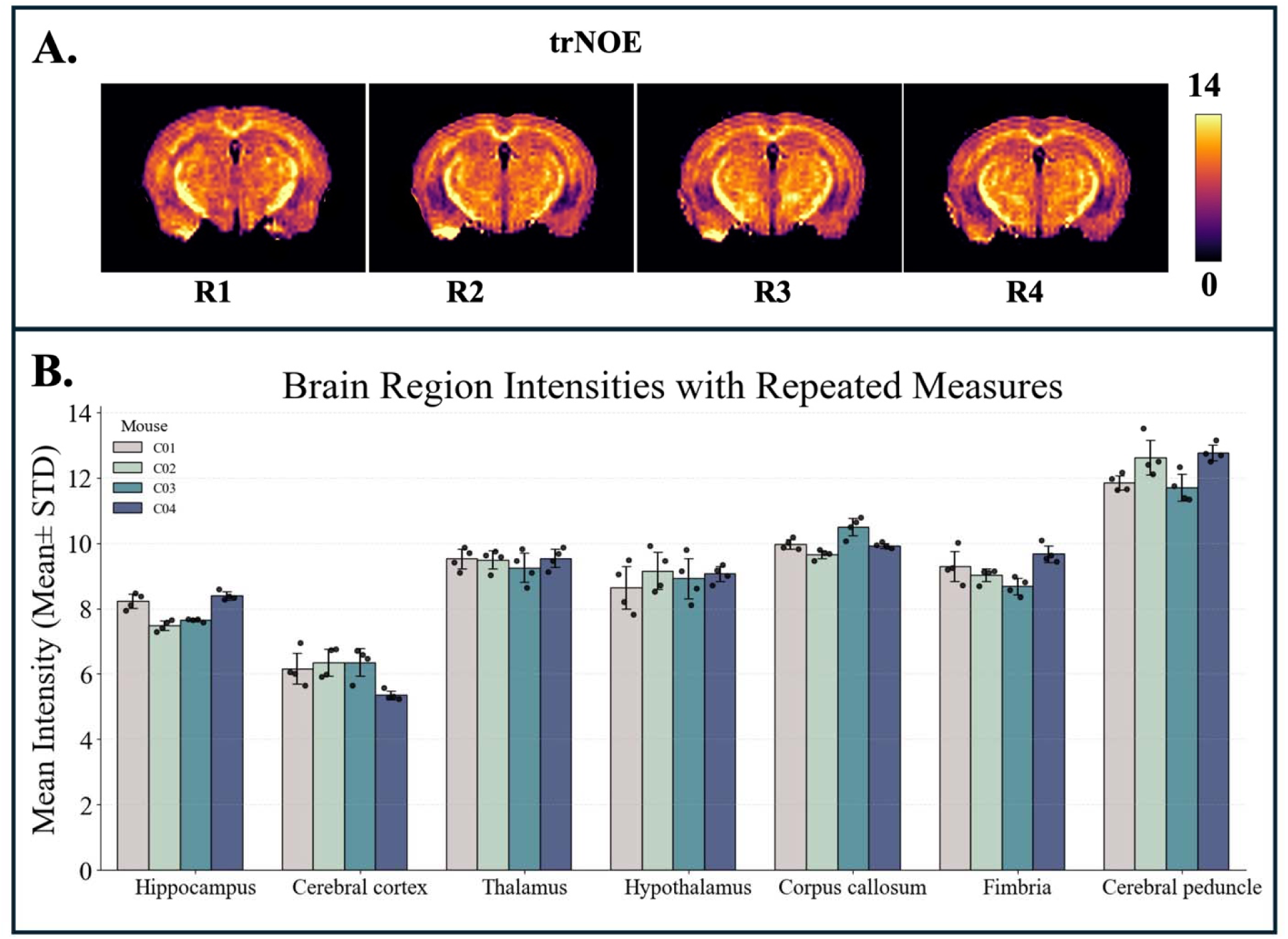
A) Representative coronal slices from four repeated sessions (R1-R4) show consistent spatial patterns in tNOE maps. B) Bar graph showing the mean tNOE intensities (mean ± standard deviation) across seven brain regions for four WT mice (C1–C4), each with four repeated measures.

### Whole brain analysis

Intra-subject COV for NOE_MTR_ ranged from 1.92% to 3.40%, while the inter-subject COV was 1.50%. The tNOE method demonstrated excellent repeatability for the whole slice, with an intra-subject COV ranging from 0.75%-5.57% and an inter-subject COV of 2.97%. MWF exhibited the highest consistency across scans, showing an intra-subject COV between 0.47% and 2.03%, and the lowest inter-subject COV of 0.75% (Fig. 5). Overall, these findings underscore the robustness of all imaging methods (NOE_MTR_, tNOE, and MWF) and their potential as reliable biomarkers for studies targeting brain lipid metabolism and lipid integrity in preclinical models.

**Figure 5:**
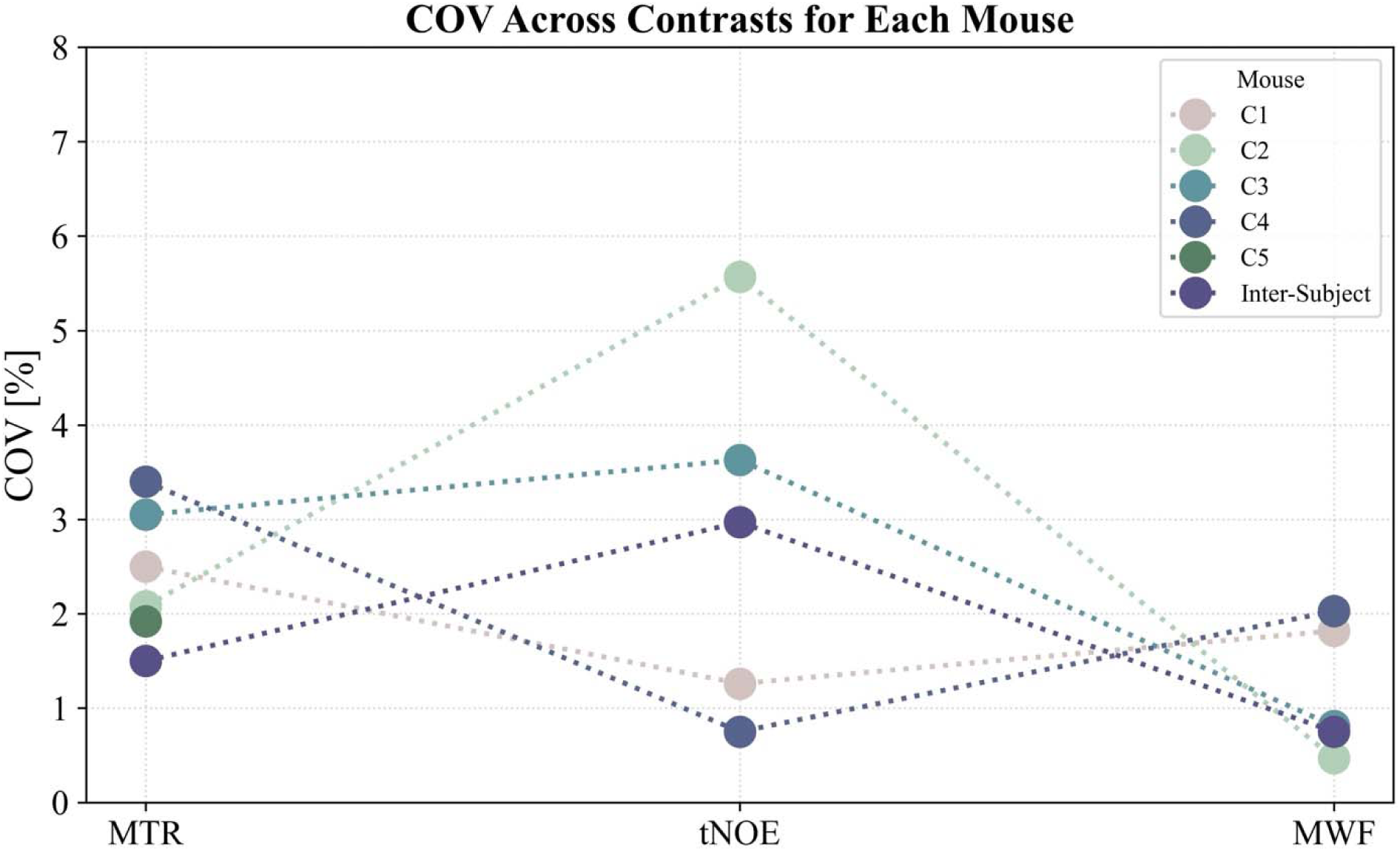
Intra-subject and inter-subject COV% for whole-brain NOE_MTR,_ tNOE, and MWF contrast maps across each mouse with four repetitions.

### Sub-regional analysis

As seen in Fig. 6, the NOE_MTR_ method demonstrated high sub-regional repeatability, with intra-subject COVs ranging from 1.21% to 7.81%. The lowest variability was observed in the thalamus and hippocampus, both showing less than 2%. In contrast, the cerebral peduncle showed higher variability in some animals. Inter-subject variability remained low across all regions, ranging from 1.82% to 4.27%.

tNOE displayed low intra-subject COVs in the majority of regions, with the thalamus and corpus callosum exhibiting COVs less than 2%. Variability was slightly increased in the fimbria and cerebral cortex. Inter-subject COVs were below 8% in all regions. MWF exhibited excellent repeatability, with intra-subject COVs ranging from 0.72% to 6.23% and inter-subject COVs varying from 0.83% to 2.78%. The most stable white matter regions were the corpus callosum and fimbria, supporting the reliability of MWF as a measure of myelin content. These findings demonstrate the robustness of NOE_MTR_, tNOE, and MWF across multiple brain regions, highlighting their potential as repeatable imaging biomarkers for quantifying lipid integrity, especially in longitudinal studies.

**Figure 6:**
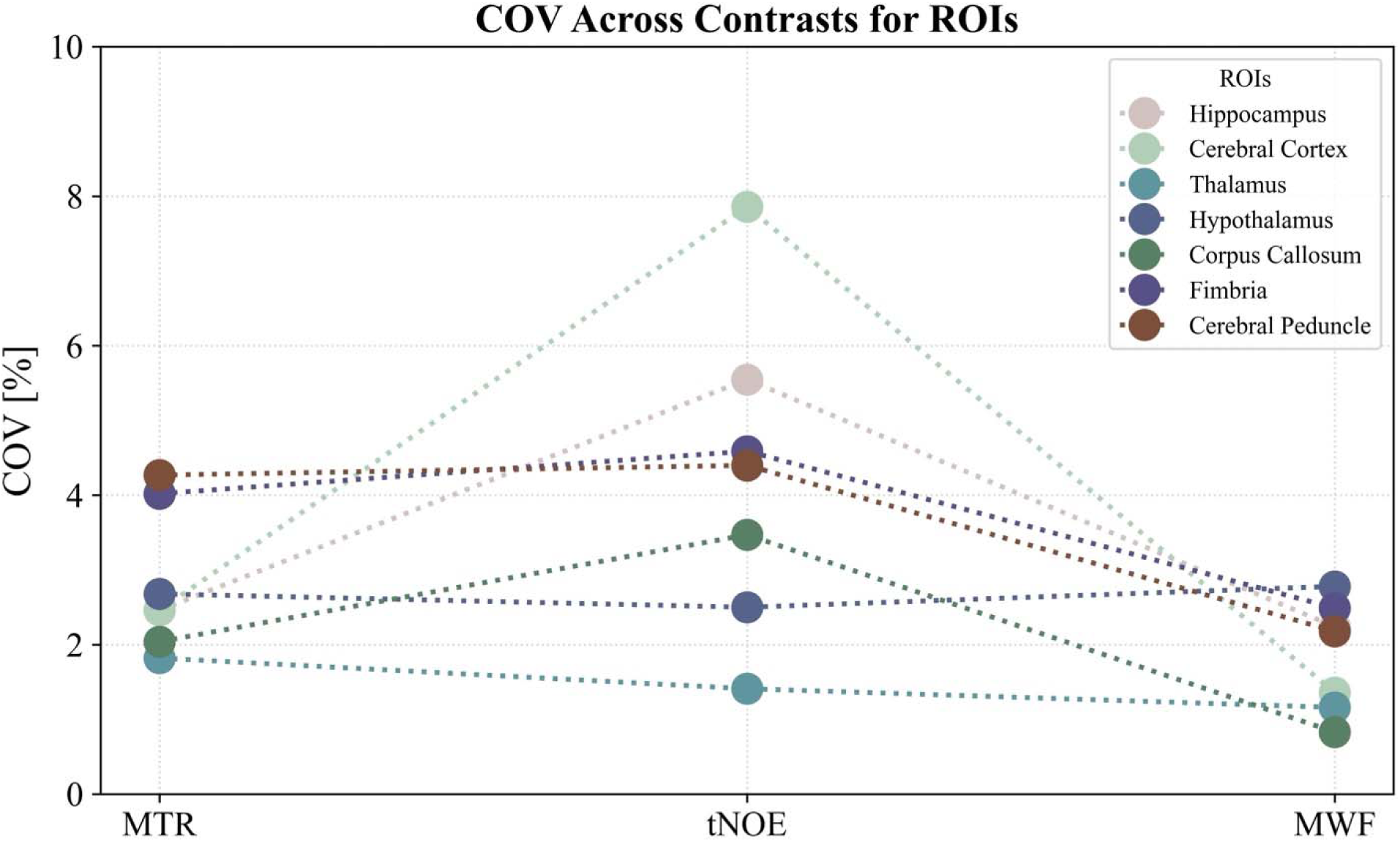
The inter-subject COV% for each ROI across various MR imaging contrast techniques, including NOE_MTR_, tNOE, and MWF map contrasts. Each cell represents the COV%, with color intensity indicating the degree of variation: blue tones denote lower variability, while red tones indicate higher variability.

### Correlation of Imaging indices with IHC

Both LFB and MBP IHC stains showed stronger staining in myelinated regions compared to non-myelinated regions (Fig. 7), as previously reported by other studies (35,36). Both qualitative and quantitative analyses revealed that tNOE and MWF imaging demonstrated a consistent association with histological markers of myelin (Fig. 7A). Specifically, tNOE values exhibited a strong positive correlation with optical density measurements from both LFB-stained sections (r = 0.72, p < 0.001) and MBP immunostaining (r = 0.83, p <0.001). This indicates that tNOE contrast is highly sensitive to myelin content (Fig. 7B). In comparison, MWF also showed a significant correlation with MBP (r = 0.63, p = 0.003), while a positive correlation with LFB did not reach significance (r = 0.34). Similarly, NOE_MTR_ showed a significant correlation with MBP (r = 0.57, p = 0.009), while its positive correlation with LFB did not reach significance (r = 0.42, p = 0.06) (Fig. 7B). These correlations were consistently observed across major white matter and gray matter regions, including the corpus callosum, fimbria, cerebral peduncle, cerebral cortex, and hippocampus. Collectively, these findings validate tNOE as a sensitive MRI biomarker of myelin integrity, with a strong association with histologically quantified myelin content.

Additionally, a strong positive correlation was observed between tNOE and MWF in white matter regions (r = 0.90, p <0.001; Supplementary Fig. 1A). Similarly, MBP and LFB levels demonstrated a significant positive correlation across various brain regions (r = 0.83, p<0.001; Supplementary Fig. 1B).

**Figure 7:**
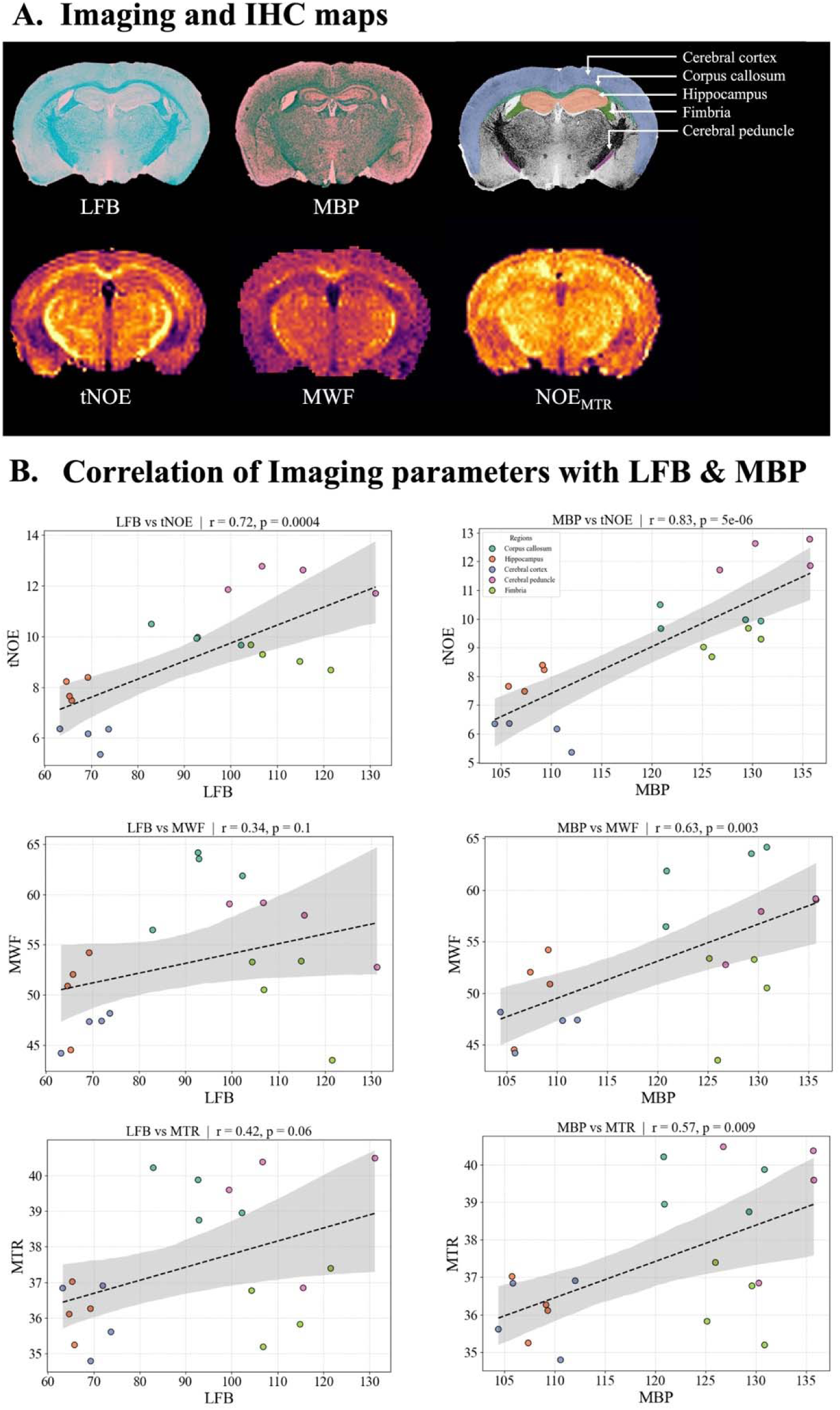
A) Representative coronal sections of the mouse brain stained with LFB and MBP demonstrate regional myelin distribution. Corresponding MRI maps of tNOE, MWF, and NOE_MTR_ show signal patterns consistent with histological staining. B) Scatter plots on the right display Pearson correlation analyses between histological optical density values, LFB and MBP, and imaging-derived metrics, tNOE, MWF and NOE_MTR_.

## DISCUSSION

We evaluated the various imaging contrast mechanisms (NOE_MTR_, tNOE, and MWF imaging) that are sensitive to lipid interactions with bulk water and systematically assessed each imaging modality for repeatability and sensitivity across the whole brain and selected ROIs. The low intra- and inter-subject COV values for whole brain and specific ROIs validate the repeatability of these methods, and correlations with IHC measures highlight the robustness of these imaging methods for investigating lipid integrity, which is crucial for preclinical and translational research on neurodegenerative diseases and other neurological disorders (10,37).

ROI analysis has highlighted that the thalamus and hippocampus emerged as the most stable anatomical structures, supporting prior reports that identified these deep brain regions as reliable sites for MR imaging contrast analysis due to reduced motion and consistent B homogeneity (38,39). Nonetheless, the overall COVs for other brain regions remained within an acceptable limit of below 8%, indicating feasibility for detecting pathological changes in lipid metabolism.

Compared to NOE_MTR_ and MWF, tNOE provided better contrast in white matter structures and demonstrated a stronger spatial correlation with immunohistochemical staining for myelin. These findings substantiate the hypothesis that tNOE imaging offers improved specificity for mobile lipid components associated with myelin (9). The frequency-selective inversion in tNOE likely accounts for its improved repeatability due to reduced confounding effects from MT and spillover. Additionally, its high repeatability suggests that tNOE has the potential to assess longitudinal biochemical changes in lipid-rich tissues with high stability and consistency. This highlights its clinical translational potential, particularly for detecting demyelination in patients with multiple sclerosis, traumatic brain injury, and neurodegenerative diseases.

MWF displayed high repeatability, with intra-subject COVs as low as 0.47% and inter-subject COVs not exceeding 3.31% across any region. These findings support the well-established position of MWF as a robust myelin biomarker in both clinical and preclinical research (40). Its stability across repeated scans and across different anatomical regions positions it as a reference standard for validating emerging contrast mechanisms like tNOE. Though MWF imaging is widely regarded as a reference standard for myelin quantification, it has inherent drawbacks that limit its sensitivity to dynamic myelin changes. MWF is an indirect measure of myelin, relying on the separation of water signal components (myelin-associated water vs. intra/extracellular water) primarily through multi-echo T_2_ relaxometry, which can be confounded by partial volume effects, exchange processes, and long acquisition times (41). In contrast, tNOE provides a more direct measure of mobile lipid components associated with myelin, as evidenced by its strong correlation with histological markers such as LFB and MBP staining. The improved biochemical specificity of tNOE, combined with its shorter scan times and better contrast-to-noise, underscores its potential as a more sensitive and translationally viable imaging biomarker for detecting subtle myelin alterations that MWF might not capture. Together, the concordance between MWF and tNOE findings supports a complementary framework in understanding myelin changes across pathological conditions. MWF may serve as a structural reference, while tNOE provides enhanced biochemical sensitivity. This dual approach allows for a more comprehensive understanding of myelin alterations.

For advanced MR imaging contrasts to be reliably adopted in clinical studies, their reproducibility must be thoroughly validated. The reproducibility crisis in biomedical imaging has highlighted the challenges associated with the lack of standardized acquisition protocols, scanner instabilities, and biological variability, all of which can impact quantitative measurements (42). This issue is particularly relevant for NOE_MTR_, tNOE, and MWF techniques, which rely on subtle changes in saturation transfer and exchange dynamics that can be easily confounded by experimental noise. Hence, assessing the repeatability of these techniques in a controlled experimental framework is critical to ensure the robustness of findings and their utility in translational research.

In this study, we utilized 2D tNOE for our proof of concept. However, acquiring 3D tNOE, which we are currently developing, is feasible and allows for more comprehensive spatial coverage and volumetric assessment. Our center has already developed and implemented a partial 3D on 7T clinical scanners and is in the process of applying these methods to evaluate age-dependent lipid changes in healthy volunteers and in patients with multiple sclerosis. Furthermore, emerging fast-acquisition strategies such as compressed sensing, parallel imaging, and magnetic resonance fingerprinting facilitate the rapid acquisition of fully sampled Z-spectra (43,44). While our findings are based on a small sample size of five WT mice, future studies will include disease cohorts to further assess the sensitivity and specificity of these methods in detecting pathological changes.

## CONCLUSION

This study demonstrates that advanced MR imaging techniques - NOE, tNOE, and MWF - provide consistent and repeatable measurements for assessing lipid and myelin content in the healthy mouse brain. The low intra- and inter-subject variability observed in both whole-brain and regional analyses supports the robustness of these methods for preclinical imaging. Notably, the tNOE imaging technique stands out for its enhanced specificity to mobile lipids and myelin-associated structures, showing strong agreement with histological markers. The findings establish a solid foundation for the broader application of these techniques to longitudinal studies and disease cohorts, especially for demyelinating and neurodegenerative disorders.

## Acknowledgements

This research was supported by the National Institutes of Health, specifically through the National Institute of Biomedical Imaging and Bioengineering (Award Number: P41EB029460) and the National Institute on Aging (Award Numbers: R01AG063869, RF1AG087306, and R01AG091760).

## Data availability

The datasets generated and/or analyzed during the current study are available from the corresponding author upon reasonable request.

## Author Contributions

S.K.K. and A.S. contributed equally to this work. S.K.K. and A.S. acquired and analyzed the data and wrote the manuscript. N.D.S., H.J., A.M., D.R., and B.B. assisted with animal preparation and imaging data acquisition. D.K. and R.P.R.N. provided critical input on the manuscript and data interpretation. M.H. and R.R. provided critical input on study design, data interpretation, and manuscript revisions. All authors reviewed and approved the final manuscript.

## Clinical trial number

not applicable.

---

**Supplementary Figure 1:**
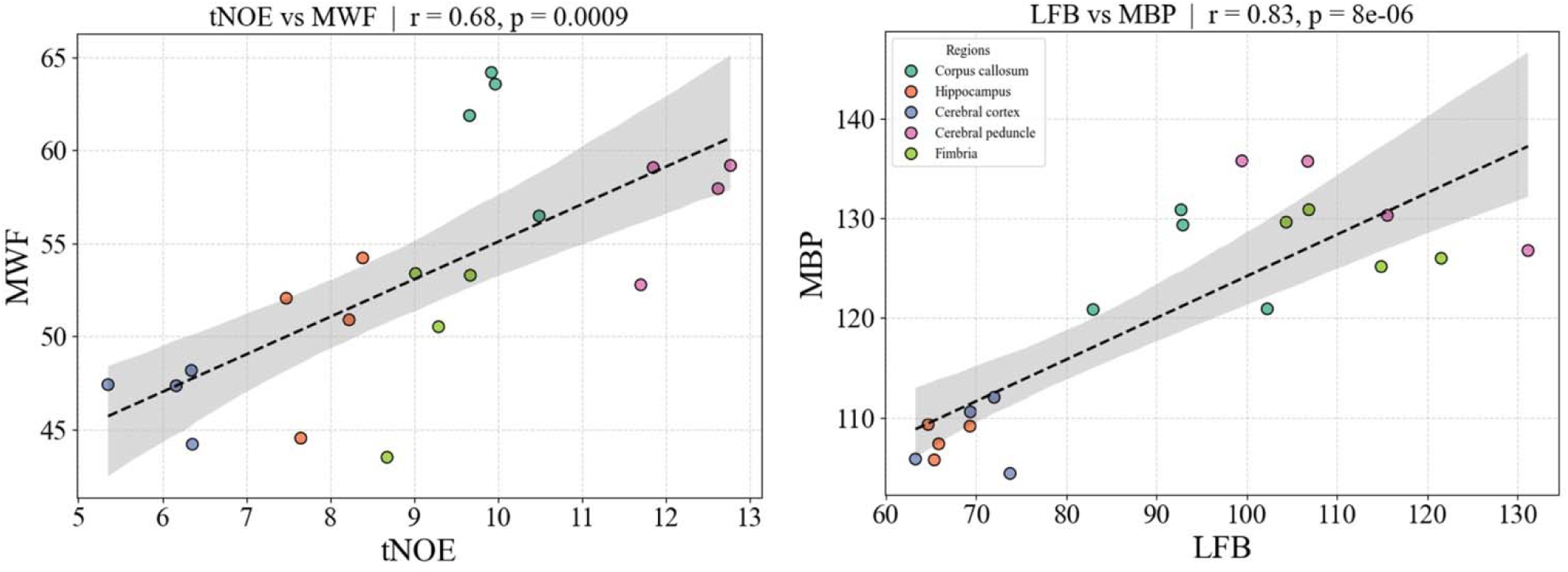
Scatter plots display Pearson correlation analyses: A. Imaging-derived metrics from white matter regions, corpus callosum, and cerebral peduncle tNOE, and MWF; B. MBP and LFB from different brain regions.

**Supplementary Table 1:**
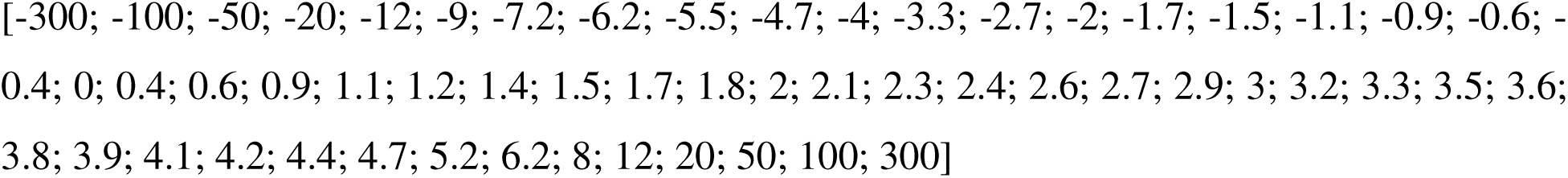
Frequency offsets used for NOE acquisition. Fifty-six frequency offsets were applied symmetrically about the water resonance to sample the Z-spectrum for NOE contrast.

**Supplementary Table 2:**
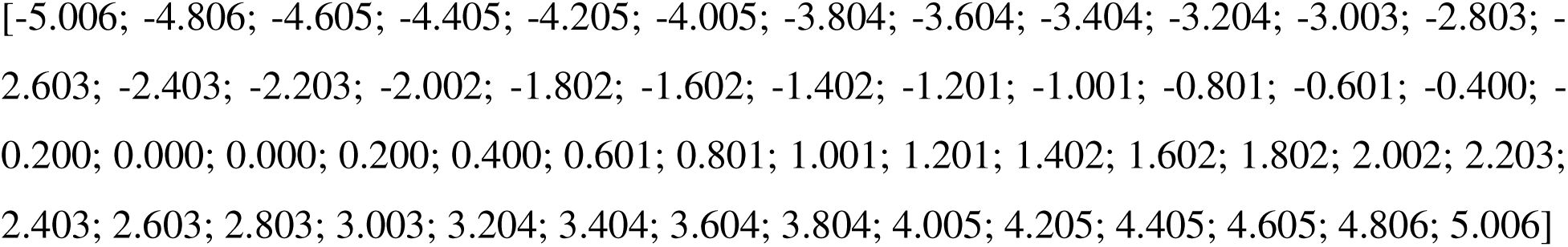
Frequency offsets used for tNOE acquisition. Fifty-two frequency offsets were applied symmetrically about the water resonance to sample the Z-spectrum for tNOE contrast.

